# A Strategic Blend of Stabilizing Polymers to Control Particle Surface Charge for Enhanced Mucus Transport and Cell Binding

**DOI:** 10.1101/2024.09.17.613453

**Authors:** Corey A. Stevens, Boris Sevarika, Brian K. Wilson, Chia-Ming Wang, Gerardo Cárcamo-Oyarce, George Degen, Timothy Kassis, Claus Michael Lehr, Rebecca Carrier, Katharina Ribbeck, Robert K. Prud’homme

**Affiliations:** Department of Biological Engineering, Massachusetts Institute of Technology, Cambridge, MA 02139; Department of Biological Engineering, Koch Institute for Integrative Cancer Research, Massachusetts Institute of Technology, Cambridge, MA 02139; Helmholtz Institute for Pharmaceutical Research Saarland (HIPS), Helmholtz Centre for Infection Research (HZI), D-66123 Saarbrücken, Germany; Department of Pharmacy, Saarland University, D-66123 Saarbrücken, Germany; Department of Chemical and Biological Engineering, Princeton University, Princeton, NJ 08544; Department of Chemical Engineering, Northeastern University, Boston, MA 02115

## Abstract

Mucus layers, viscoelastic gels abundant in anionic mucin glycoproteins, obstruct therapeutic delivery across all mucosal surfaces. We found that strongly positively charged nanoparticles (NPs) rapidly adsorb a mucin protein corona in mucus, impeding cell binding and uptake. To overcome this, we developed mucus-evading, cell-adhesive (MECS) NPs with variable surface charge using Flash NanoPrecipitation, by blending a neutral poly(ethylene glycol) (PEG) corona for mucus transport with a small amount, 5 wt%, of polycationic dimethylaminoethyl methacrylate (PDMAEMA) for increased cell targeting. *In vitro* experiments confirmed rapid mucus penetration and binding to epithelial cells by MECS NPs, suggesting a breakthrough in mucosal drug delivery.

## Introduction

The effective delivery of therapeutics is often hindered by biological barriers, particularly mucus layers found throughout the body’s mucosal surfaces^1,2^. Mucus layers represent a significant barrier to therapeutic drug delivery due to their complex structure and interactions with drug carriers^3,4^. Ideally, delivery systems should navigate through mucus while still effectively binding to target cells, a balance that has proven difficult to achieve^5^.

To address this critical need, we developed mucin-evading cell-adhesive nanoparticles (MECS NPs). Our NP design, incorporating these seemingly contrary properties, was informed by examining how mucus interacts with particles and affects their mucosal drug delivery performance. We observed that strongly positively charged NPs form a protein corona within mucus, which inhibits cellular interactions. To address this, we engineered NPs using a blend of positively charged (PDMAEMA) and neutral (PEG) polymers. PDMAEMA facilitates cellular binding and uptake (“targeting”), while PEG enables muco-penetration and prevents mucoadhesion. A rapid precipitation process, Flash NanoPrecipitation, enables the assembly of NPs with dense polymer coronas, but variable surface charge by altering the ratio of neutral to charged stabilizers. By overcoming the long-standing challenge of providing both mucus transport and epithelial cell targeting, our newly developed NPs have the potential to significantly advance mucosal drug delivery.

## Results and Discussion

### Positively charged, electrostatically stabilized latex NPs acquire a protein corona in mucin

Positively charged NPs have garnered significant interest in the field of drug delivery because of their potential for cellular targeting and uptake^6^. The electrostatic attraction between positively charged particles and negatively charged cell membranes facilitates their interaction and subsequent cellular uptake. However, successful drug delivery through mucosal surfaces necessitates traversing the mucus gel layer, which contains negatively charged mucin polymers. Mucins, the main gelling component of mucus, are heavily glycosylated polyanionic proteins that cross-link through a series of covalent and non-covalent bonds, resulting in a mesh-like network^7^. While a positive charge can promote cellular targeting by NPs, traditionally, the interaction between positively charged particles and the negatively charged mucins has been assumed to lead to significant adhesion and reduced diffusion^8^.

To investigate the diffusion of NPs within the mucus barrier, we employed a well-defined model system composed of purified mucins from the intestine (MUC2) and lungs/stomach (MUC5AC) that recapitulates the properties of native mucus (Fig. 1a–c, Supplementary Fig. S1). Using single-particle tracking (SPT), we quantified the diffusion of positively and negatively charged, electrostatically stabilized polystyrene latex NPs of various sizes (100 nm, 200 nm, 500 nm, and 1000 nm) within the mucin gels. As a control, we tracked NPs diffusing within a water–glycerol mixture, and the NPs exhibited the expected inverse correlation between particle size and diffusivity (Fig. 1d). Intriguingly, the diffusivity of positively charged NPs in the mucin gels was not significantly different from that of negatively charged NPs of the same size (Fig. 1e [MUC2], 1f [MUC5AC]). This observation expands the traditional understanding of mucoadhesion for positively charged NPs^8,9^.

**Figure 1.**
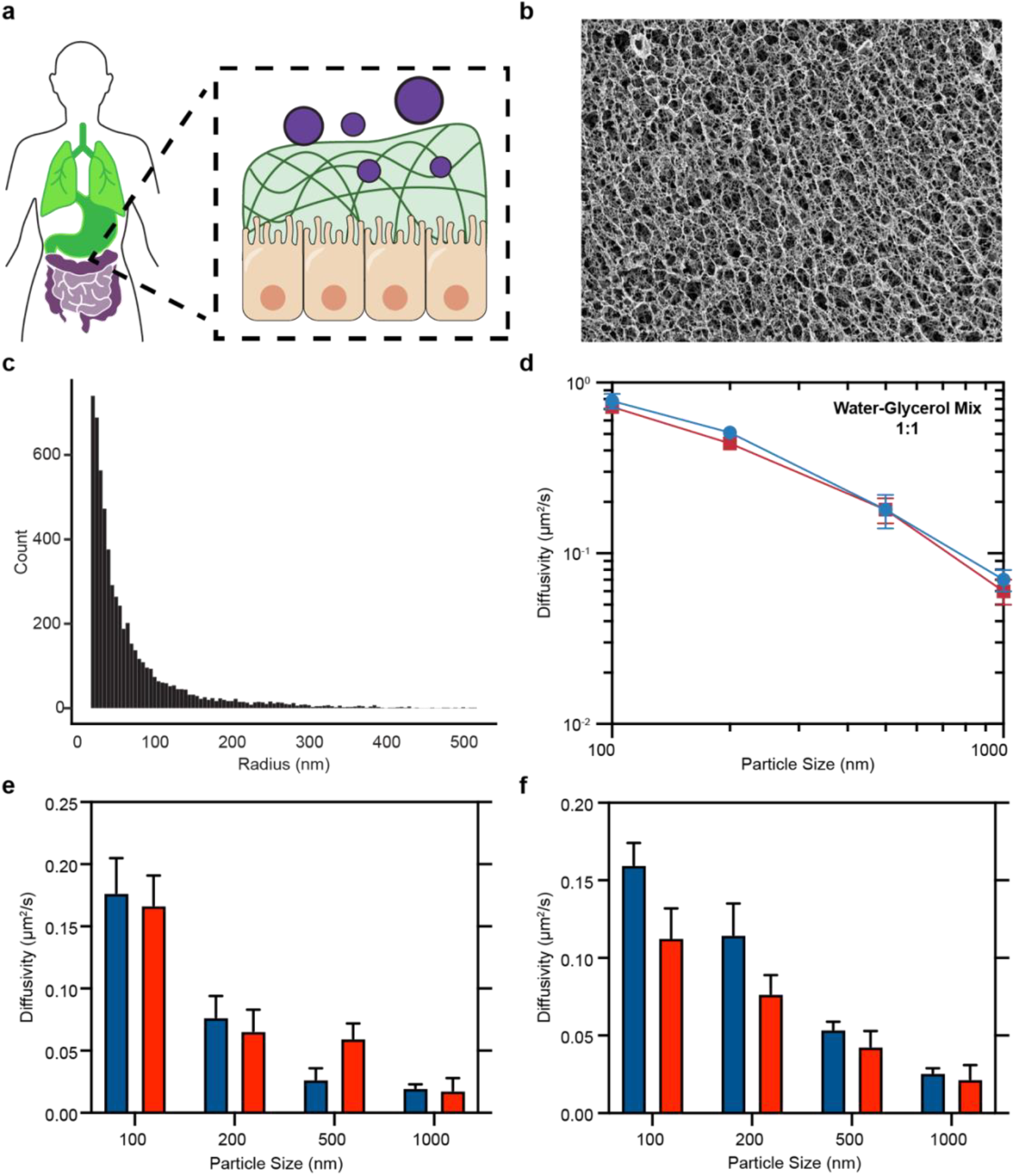
NP transport across the mucus barrier. (a) Schematic representation of mucosal tissue and NPs for drug delivery across the mesh-like mucus network. (b) CryoSEM image of a reconstituted 1 wt% MUC2 gel. (c) Two-dimensional pore analysis of cryoSEM images of 1 wt% MUC2 gels. Counts represent the number of pores with the corresponding radius. (d) SPT-determined diffusivity of polystyrene NPs in a 1:1 water:glycerol mixture (anionic particles in blue, cationic particles in red). (e–f) Diffusivity of positively (red) and negatively (blue) charged NPs in 1 wt% (e) MUC2 and (f) MUC5AC gels.

To further explore the interaction between mucin gels and NPs, we imaged NPs within mucin gels by cryo-scanning electron microscopy (cryoSEM). Our imaging reveals distinct interactions based on particle charge. Negatively charged NPs have minimal interaction with mucin chains, with strands seemingly draped across the particle surface (Fig. 2a). In contrast, positively charged NPs are completely enveloped by a dense mucin layer (Fig. 2b). These observations suggest the formation of a “mucin corona,” i.e., a coating of mucins adhered to the surface of cationic NPs. This observation mirrors the well-documented protein corona; where proteins in biological fluids adsorb onto particle surfaces, altering their properties and influencing interactions with the surrounding environment^10–12^. Zeta potential measurements show charge reversal; cationic NPs transition from positive to negative after mucin exposure, which is consistent with their diffusive behavior (Fig. 2c, Supplementary Table S1). The mucin corona masks the charge of the cationic NPs, making them anionic, causing them to diffuse like anionic NPs (Fig. 2d–e)^13^. Thus, cationic NPs are not significantly trapped within the mucin gel; instead, they become coated in mucin, which may significantly impact the behavior and fate of NPs within mucus.

**Figure 2.**
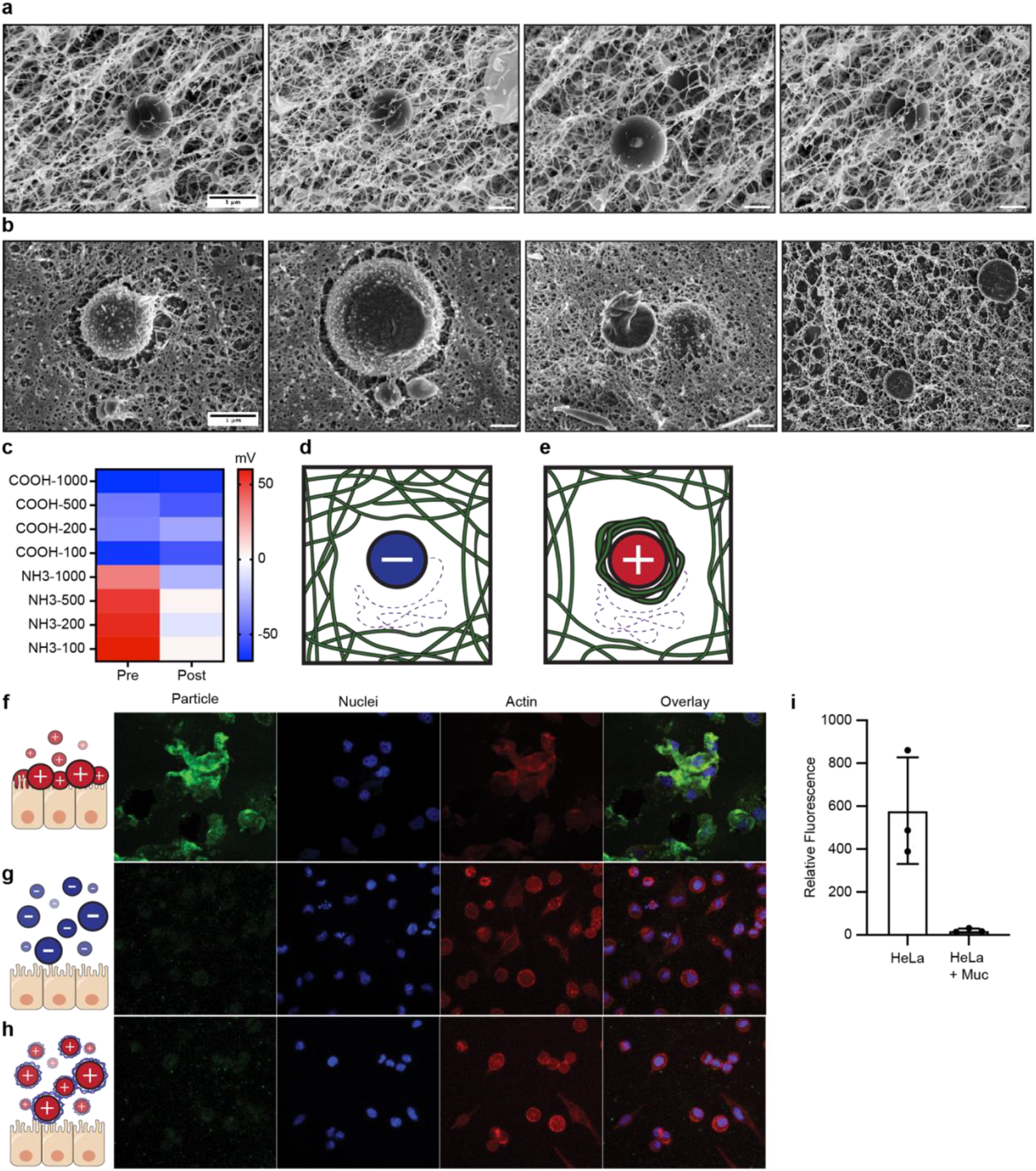
Formation of a mucin corona on cationic NPs. (a–b) CryoSEM images of 1 μm (a) anionic and (b) cationic particles in 1 wt% MUC2 gels. (c) Zeta potential measurements of particles pre- and post-incubation with MUC2. (d) Schematic representation of anionic NPs diffusing in mucin gel. (e) Schematic representation of cationic NPs with a mucin corona diffusing in a mucin gel. (f–h) Confocal images depicting the influence of the mucin corona on NP adhesion to epithelial cells: (f) cationic NPs adhering to HeLa cell monolayers, (g) anionic NPs incubated with HeLa cell monolayers, and (h) cationic NPs pretreated with MUC2 prior to incubation with a HeLa cell monolayer (green: particles, blue: nuclei [DAPI], red: actin [phalloidin]). (i) Quantification of microscopy images shown in f–h.

### Mucin coronas hinder cellular targeting

To explore the impact of the mucin corona on cell targeting, we investigated particle adhesion to epithelial cells. We grew a monolayer of HeLa cells overlaid with MUC2 (100 μL of 1 wt% MUC2; Fig. 2f–h). In the absence of mucin, cationic NPs electrostatically adhere to the negatively charged epithelial cell surface (Fig. 2f), as evidenced by green NP fluorescence colocalizing with nucleus and actin fluorescence. Conversely, anionic NPs do not accumulate on the epithelial cell layers (Fig. 2g), consistent with established principles in drug delivery^14^. Crucially, cationic NPs pre-incubated with mucus no longer exhibit electrostatic adhesion to the surface of epithelial cell monolayers due to the mucin corona (Fig. 2h–i).

Our data demonstrate that while a positive charge aids in cellular targeting, it also leads to the accumulation of a mucin corona within mucosal tissues, thereby impeding targeting and potentially inhibiting drug release. This underscores the need for innovative NP designs.

### Library of MECS NPs shows tunable mobility through mucins

To design NPs with enhanced mucosal drug delivery capabilities, we designed MECS NPs that combine the cellular targeting benefits of a positive charge with the transport-promoting properties of PEG^15,16^. PEGylation not only reduces the overall positive charge density on the particle surface but also sterically hinders extensive adherence by mucins, potentially preventing the formation of a thick mucin corona.

To produce MECS NPs, we used Flash NanoPrecipitation, a kinetically controlled, block-copolymer-directed assembly process (Fig. 3a)^17,18^. The resulting NPs consist of a core composed of hydrophobic poly(styrene) homopolymer, mixed with the adsorbed hydrophobic portion of the stabilizing polymer, and a dense hydrophilic polymer brush corona surrounding the NP, as depicted in Fig. 3b. The charge of this polymer corona, crucial for targeting, can be tailored by adjusting the ratio of neutral (polystyrene-block-PEG), negatively charged (poly(acrylic acid) [PAA]), or positively charged (PDMAEMA) chains in the stabilizing layer (Fig. 3c–e).

**Figure 3.**
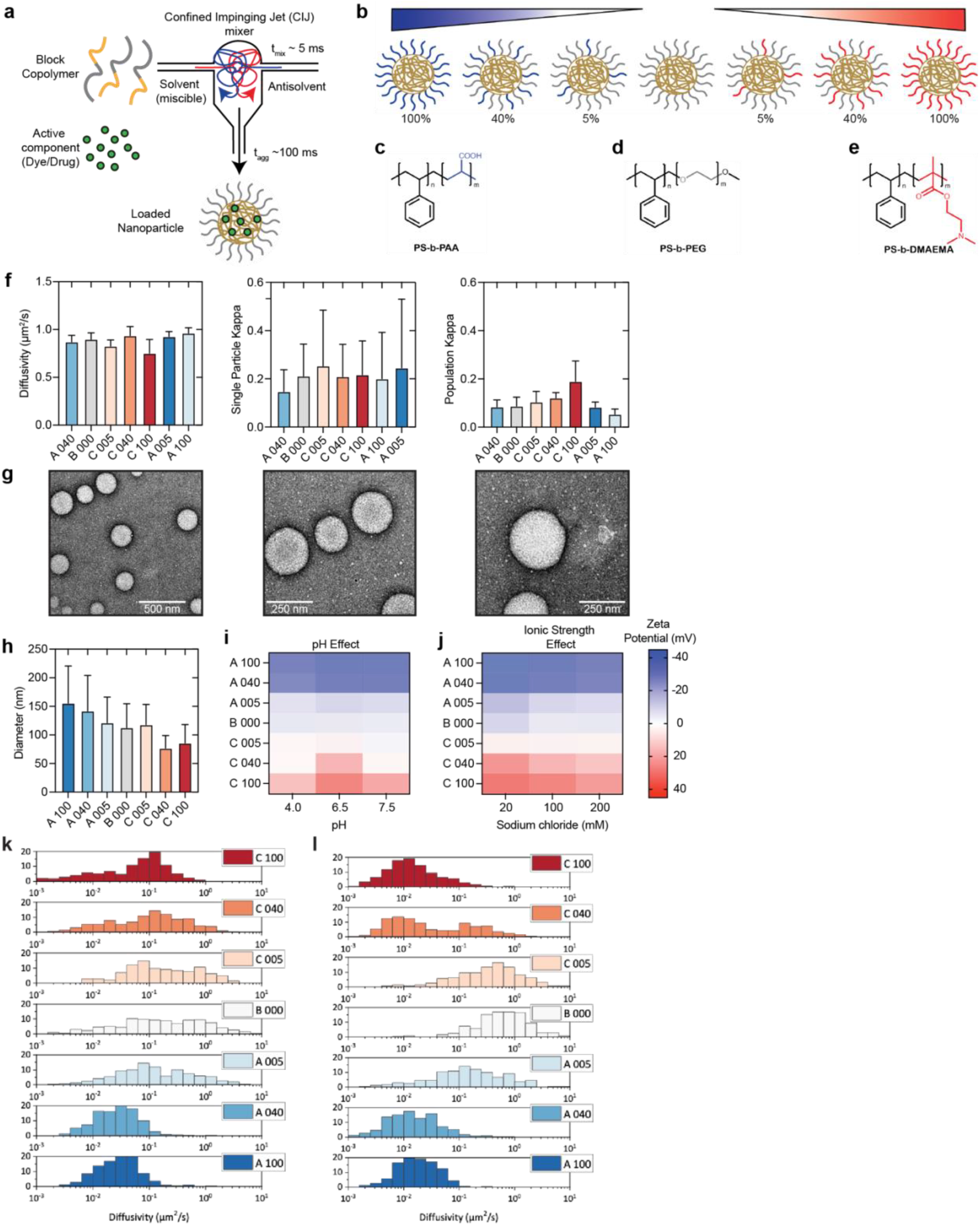
Production of MECS NPs by Flash NanoPrecipitation. (a) Schematic representation of the Flash NanoPrecipitation procedure used to produce an MECS NP library. (b) Representation of NPs tested for mucosal transport and cell targeting. (c–e) Diagrams of (c) negatively charged PAA, (d) polystyrene-block-PEG, and (e) positively charged PDMAEMA. (f) SPT-determined diffusivity of NPs produced by Flash NanoPrecipitation in a 1:1 water:glycerol mixture (left), with individual particle kappa values (middle) and kappa values averaged over the entire population of tracked particles (right). (g) Transmission electron microscopy images of B000 particles, showing polymers brushed about the particle surfaces. (h) Dynamic light scattering size measurement of each NP. (i–j) Zeta potential measurements of each particle in solutions of different (i) pH and (j) ionic strength. (k–l) Histogram showing the diffusivity of every tracked particle in 1 wt% (k) MUC2 or (l) MUC5AC.

We characterized these NPs by transmission electron microscopy, SPT, dynamic light scattering, and zeta potential measurements (Fig. 3f–j). The NPs in our library are approximately 100 nm in diameter and exhibit variable and controlled surface charges, ranging from -38 mV to +40 mV depending on the ratio of PAA:PEG or DMAEMA:PEG stabilizer block copolymers (Fig. 3h–j). Low levels of polyelectrolyte stabilizer addition, C005 and A005, yield NPs with nearly neutral surfaces (|ζ| < 10 mV). All NPs were loaded with the fluorescent dye Hostasol Yellow 3G at a concentration of 2 wt%, to facilitate imaging and to mimic encapsulation of a hydrophobic drug. Importantly, the hydrophobic dye is confined within the NP core, eliminating any potential interference with mucous interactions, which can occur with surface-functionalized dyes.

We investigated the diffusion behavior of seven MECS NP types (A100, A040, A005, B000, C005, C040, C100) in reconstituted MUC2 and MUC5AC gels by SPT. As a control, we measured NP diffusion in polyanionic synthetic carboxymethyl cellulose (CMC) hydrogels (Supplementary Fig. S2–S4). As expected, neutral particles coated only with PEG (B000) exhibited high mobility in mucin gels. Conversely, particles with high PDMAEMA content (C100, C040) or high PAA content (A100, A040) displayed lower mobility in both mucin gels. Interestingly, our data show that a minimal amount (5%) of PDMAEMA (C005) resulted in particles with diffusion properties comparable to those of fully PEGylated particles in MUC2 and MUC5AC gels (Fig. 3k [MUC2], 3l [MUC5AC]). These polymer-stabilized NPs present a hydrophilic polymer brush independent of the amount or charge of added polyelectrolyte block copolymer, unlike the electrostatically stabilized latex NPs characterized in Fig. 1–2 that present a hydrophobic poly(styrene) surface between singly charged functional groups. This observation hints at the potential for these particles to achieve both efficient muco-transport and targeted epithelial cell interactions.

### MECS NPs achieve both mucus transport and epithelial cell targeting

To determine if 5% PDMAEMA (C005) particles target to epithelial cells, we conducted *in vitro* cell culture experiments. We grew epithelial cell (HeLa) monolayers and overlaid a 1 wt% mucin gel (MUC2). Subsequently, we added C005 particles on top of the mucin layer and incubated for 1 h; for controls, we prepared similar samples with A005 and B000 particles.

After thorough washing and fixation, we stained actin (phalloidin) and nuclei (4’,6-diamidino-2-phenylindole [DAPI]). Confocal microscopy revealed minimal colocalization of green fluorescent NPs with DAPI and phalloidin in the A005 and B000 conditions (Fig. 4a–b). Conversely, the C005 particles displayed extensive colocalization with both nuclei and actin filaments, indicating successful adhesion to epithelial cells (Fig. 4c). Flow cytometry results further confirmed this observation, demonstrating significantly higher green fluorescence intensity in cells exposed to C005 particles compared with those treated with B000 or A005 particles (Fig. 4d). Together, these results strongly suggest that the combination of PEG for mucus transport and small amounts of PDMAEMA for positive charge enables these MECS NPs to achieve both efficient mucus penetration yet still show increased epithelial cell targeting.

**Figure 4.**
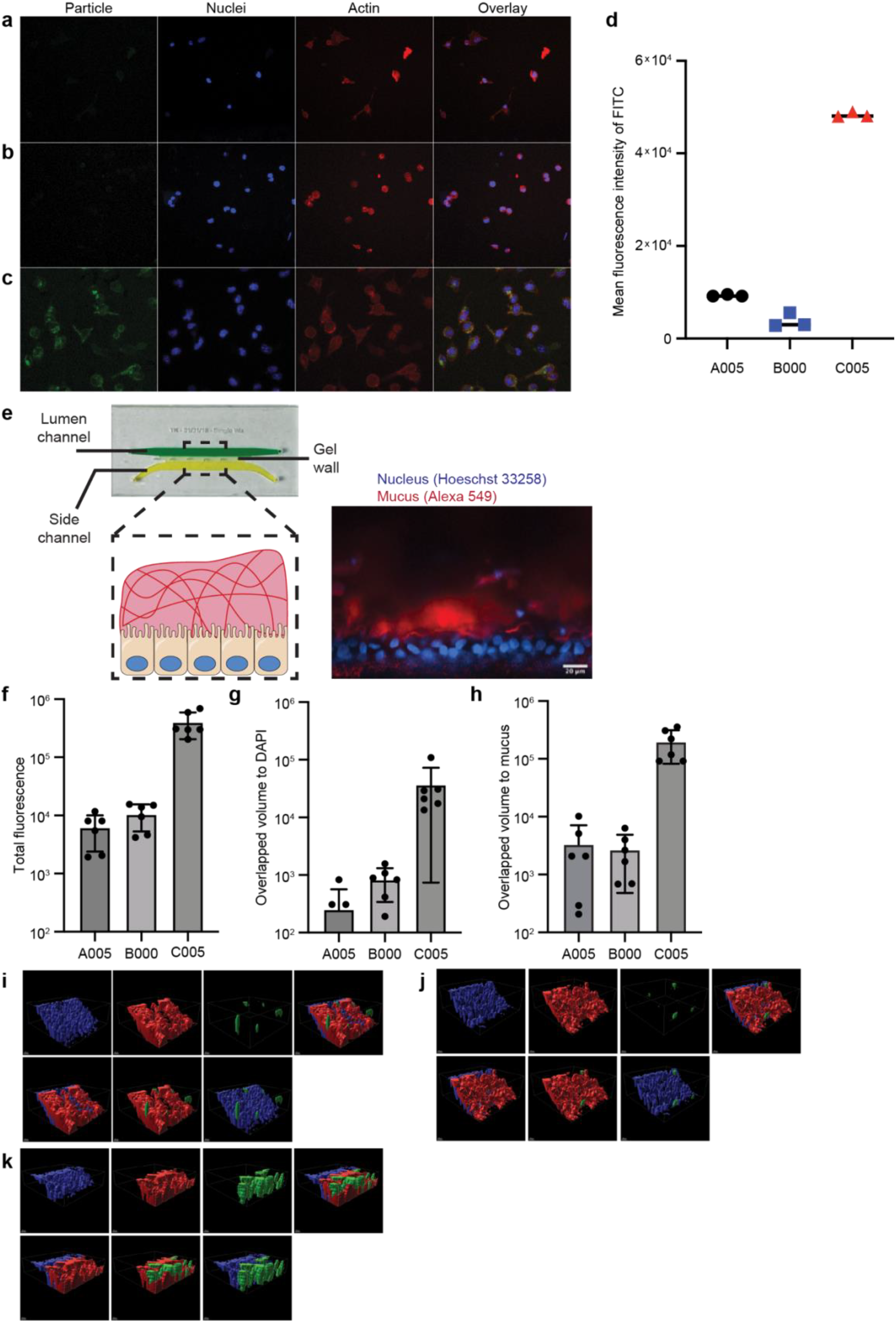
Superior epithelial cell targeting of MECS NPs. (a–c) Confocal microscopy images of HeLa cells overlaid with 100 μL of 1 wt% MUC2 and A005 (a), B000 (b), or C005 (c). (d) Flow cytometry quantification of HeLa cells coated with MUC2 and incubated with NPs. (e) Schematic representation of the mesofluidic intestinal chip (right) and an image of the mucosal interface within the system (left). (f–h) Quantification of confocal images of organoid systems incubated with equivalent amounts of A005, B000, and C005 showing the (f) total remaining fluorescence, (g) NP green fluorescence overlapped with nuclei (DAPI), and (h) NP green fluorescence overlapped with mucus (WGA-Texas Red). (i–k) Volumetric analysis of (i) A005, (j) B000, and (k) C005 particles incubated with gut-on-a-chip organoid. Z-stack confocal images were converted into volumes for each fluorescent channel (blue: Hoechst; red: WGA-Texas Red; green: NPs). Hostasol Yellow 3G.

To further solidify these findings, we used a gut organoid-based intestinal model that offers a controlled environment incorporating mucus-producing goblet cells to recapitulate the *in vivo* mucosal interface^19^. Here, intestinal stem cells are seeded upon a gel matrix within a mesofluidic chip, and factors are introduced to stimulate differentiation and mucus production in a neighboring channel, mimicking the gut lumen (Fig. 4e). NPs added to the luminal channel can traverse the mucus layer and interact with underlying epithelial cells. We labeled cell nuclei (Hoechst) and mucus (wheat germ agglutinin [WGA]-Texas Red), followed by incubation with green fluorescent NPs. Confocal microscopy analysis revealed a remarkable difference in colocalization among the three tested NPs (A005, B000, C005) (Fig. 4f–k). Notably, the signal from C005 particles overlapped substantially with the DAPI channel (nuclei), indicating a five-fold increase in colocalization compared with PAA (A005) or PEG (B000) particles (Fig. 4f–h). The enhanced colocalization achieved with 5% PDMAEMA particles (∼1.5 mV) in both mucin gels and a gut organoid system indicates significantly higher cell targeting, opening exciting avenues to revolutionize mucosal drug delivery.

## Conclusion

In conclusion, mucus layers present a formidable obstacle to drug delivery, with positively charged NPs developing a mucin corona, impeding cell targeting. To overcome this, we have developed MECS NPs, blending a neutral polymer for mucus penetration with a polycationic component for increased cell targeting. Our MECS NPs demonstrate mucus penetration while maintaining cell binding interactions, marking a breakthrough in mucosal drug delivery. The effect is observed over a narrow range of compositions with 5% cationic polymer and zeta potentials of approximately 1.5 mV. These findings offer the prospect of more efficient drug delivery systems, potentially reducing required dosages and minimizing off-target effects. Moreover, this optimized delivery strategy shows potential for mucosal vaccines and addressing challenges related to mucus-associated diseases like cystic fibrosis and chronic inflammatory bowel disease.

## Author Contributions

C.A.S, C.M.L, KR and R.K.P conceived of the project. B.K.W synthesized and characterized the particles as well as carried out experiments. C.A.S and BS performed single-particle tracking experiments and data analysis. BS and G.D.D performed micro- and macro-rheological experiments. C.A.S, C.M.W and G.C.O performed gut-on-a-chip organoid experiments and analysis. TK, C.M.W and RC designed, built and characterized the gut-on-a-chip organoid system. All authors contributed to the writing and editing of the manuscript.

## Acknowledgements

This work was supported by the Bill and Melinda Gates Foundation (award no. 6948010). C.A.S was supported by the Canadian Institute of Health Research (CIHR) postdoctoral fellowship (MFE-187894). BS was supported by the DAAD (German Academic Exchange Service) Annual Scholarship for Study Abroad for Master Students 2021/22 (57555903). G.D. D was funded by NIH Training Grant # T32-ES007020. We thank Prof. Linda Griffith for her help with the gut-on-a-chip organoid design and construction. We thank the Koch Institute’s Robert A. Swanson (1969) Biotechnology Center for technical support, specifically The Peterson (1957) Nanotechnology Materials Core Facility, David Mankus and Abigail Lytton-Jean for assistance with CryoSEM imaging.

