## supplemental materials for "A Strategic Blend of Stabilizing Polymers to Control Particle Surface Charge for Enhanced Mucus Transport and Cell Binding"

### 1.1 Materials

We purchased the following materials from Sigma Aldrich, including phosphate-buffered saline (PBS) buffer, microscopy slides, coverslips, Dulbecco's modified Eagle medium (DMEM), culture flasks, and 96-well plates. Fluorescent probes (4',6-diamidino-2-phenylindole [DAPI] and phalloidin) were obtained from ThermoFisher. We purchased fluorescent polystyrene nanoparticles (NPs) from Magsphere, Inc., including negatively charged (carboxylated) NPs with a diameter of 100 nm (Catalog No. CA100NM), 200 nm (Catalog No. CA200NM), 500 nm (Catalog No. CA500NM), or 1 µm (Catalog No. CA001UM) and positively charged (amine) NPs with a diameter of 100 nm (Catalog No. AM100NM), 200 nm (Catalog No. AM200NM), 500 nm (Catalog No. Am500NM), or 1 µm (Catalog No. AM001UM).

### 1.2 ZETA POTENTIAL

We measured the zeta potential of the NPs using a Malvern Zetasizer Pro in a buffer of 1X PBS before or after coating with mucin. Here, we report the zeta potential as an average of three replicates with the corresponding sample-based standard deviation.

### 1.3 MUCIN PURIFICATION AND HANDLING

Mucin preparation followed previously established protocols[1–3]. Briefly, we purified MUC2 from fresh pig intestinal scrapings, scraped in lab from intestines collected from a local slaughterhouse. We solubilized the isolated mucus layer in a sodium chloride buffer containing sodium azide and protease inhibitors. After centrifugation to remove insoluble material, mucins were isolated via gel filtration chromatography on a Sepharose CL2B column. Finally, the isolated mucin was concentrated, desalted, lyophilized, and stored at -20°C. Before use, weighed mucin was resuspended in 1X PBS buffer to a desired concentration (wt%) and vigorously shaken at 4°C overnight.

### 1.4 NP PRODUCTION BY FLASH NANOPRECIPITATION

We prepared NPs using Flash NanoPrecipitation in a confined impinging jet mixer as previously described[4–6]. Blending neutral (polystyrene-block [PS-b]-poly(ethylene glycol) [PEG]) and charged polyelectrolyte (PS-b-poly(acrylic acid) [PAA] or PS-b-dimethylaminoethyl methacrylate [DMAEMA]) block copolymer stabilizers enables the production of NPs with continuously tunable zeta potentials between -40 mV to +40 mV. We prepared NPs by rapidly mixing a tetrahydrofuran (THF) stream containing PS homopolymer and PS-b-PEG combined with PS-b-PAA or PS-b-DMAEMA against an aqueous antisolvent of deionized water. The composition of this THF solvent stream determines the NP composition, and all NPs studied here were produced from a 5-mg/mL PS homopolymer, 5-mg/mL combined block copolymer stream. We also added Hostasol Yellow 3G at 0.1 mg/mL as a hydrophobic, fluorescent dye with similar optical properties as fluorescein isothiocyanate. Samples are given a name code, where the first letter identifies the type of polyelectrolyte block copolymer (A for PS-b-PAA, anionic; C for PS-b-DMAEMA, cationic), and the three digits indicate the weight percentage of the polyelectrolyte block copolymer in the total mass of block copolymer. For example, B000 indicates neutral PS-b-PEG-only NPs with no polyelectrolyte, and A005 indicates 5% PS-b-PAA, 95% PS-b-PEG. For A005, NPs were made from a THF solution containing 5 mg/mL PS, 0.1 mg/mL Hostasol Yellow 3G, 0.25 mg/mL PS-b-PAA, and 4.75 mg/mL PS-b-PEG.

#### 1.5 Single Particle Tracking (SPT)

Polystyrene NPs were vortexed (1 min) and then underwent a two-step dilution (1:20 and 1:60 with water) for a final concentration of 1:1200. Following sonication (5 min) to disperse aggregates, we prepared samples by combining mucin solutions or a glycerol-water mixture with the diluted NPs (30:1 volume ratio). We constructed a custom flow cell using double-sided tape to create a rectangular channel on a microscope slide with approximate dimensions of 4 mm × 25 mm × 0.2 mm, sealed with a coverslip. Approximately 20 µL of the sample was loaded and sealed with Valap to prevent evaporation. We performed SPT video acquisition on an Axio Observer D.1 microscope with a 100x objective and Hamamatsu camera (binning 2×2 for an improved signal-to-noise ratio), capturing videos at 30.3 frame/s for 10 s at room temperature within the central channel region to avoid potential wall effects. We prepared three independent samples per condition, with ten videos captured at different locations within each sample to maximize the number of analyzed fields of view.

#### 1.6 SPT Data Analysis

We analyzed images using a customized version of publicly available MATLAB codes (Furst E. M. and Pelletier V. and Kilfoil M.). SPT analysis was performed as previously described[7]. Briefly, custom MATLAB code identified and tracked particles in microscopy images based on intensity thresholds and size. Trajectories were built by linking particle positions based on spatial and temporal proximity. A maximum linking distance was chosen based on particle size and expected Brownian motion. We applied drift correction to account for unintentional microscope stage movements or bulk fluid flow. We calculated the mean square displacement (MSD) to determine diffusion coefficients (normal, sub-diffusive, or super-diffusive).

#### 1.7 Rheological Measurements

We measured the rheological properties of MUC5AC, MUC2, and carboxymethyl cellulose (CMC) solutions (0.5%, 1.0%, and 2.0%) in a HEPES buffer (20 mM) at a pH of 6.5 and a sodium chloride concentration of 100 mM using an ARESG2 rheometer (TA Instruments, New Castle, DE, USA) with a parallel-plate geometry (25 mm). The temperature was maintained at 25°C during all experiments. We performed strain-sweep measurements (from 0.1% to 100%) at a constant angular frequency of 5 rad/s. Frequency-sweep experiments were performed at an oscillation amplitude within the linear response region over a frequency range of 0.1–100 rad/s. We conducted flow-sweep measurements at a shear rate ranging from 1.0 to 100.0 $s^{-1}$. All experiments were repeated as three independent replicates, and the average value and standard deviations are reported.

#### 1.8 Cryogenic Scanning Electron Microscopy

We subjected image samples to high-pressure freezing using a specialized apparatus designed for cryo-preservation. This process involves rapidly freezing the samples under high pressure to minimize ice crystal formation and preserve their native structure. We carefully fractured the high-pressure-frozen samples while still in their frozen state using a freeze-fracture apparatus. This step created a fractured surface that exposed the inner structures of the samples. The fractured surfaces of the samples were subjected to sublimation, a controlled process in which ice is directly converted into water vapor without passing through a liquid phase. We conducted this sublimation step under carefully controlled conditions to remove any remaining ice from the fractured surfaces. Following sublimation, the samples were sputter-coated with a thin layer of platinum and carbon. The prepared samples were transferred via a Leice VCT 500 cryo-transfer system that allows for the transfer of cryo samples under vacuum and were then loaded into and imaged in a Zeiss Crossbeam 540 focused ion beam-scanning electron microscope (FIB-SEM). We obtained sample images at -130°C (5 kV, 9.4-mm working distance, 20,000x magnification) on the Zeiss Crossbeam 540 FIB-SEM using inline and secondary electrons secondary ions detectors.

### 1.9 EPITHELIAL CELL ADHESION AND QUANTIFICATION

We cultured HeLa cells using established protocols in DMEM with 10% fetal bovine serum and penicillin-streptomycin. Approximately two million HeLa cells from a frozen stock were first seeded into a T75 flask until a confluent monolayer formed. Following adherence and monolayer formation, cells were washed with 1X PBS and incubated with trypsin for 4 min. Cells were washed and incubated with NP suspensions at the chosen concentration for a specified incubation period. After incubation, we removed non-adherent NPs through washing steps and quantified adherent NPs using appropriate methods such as fluorescence microscopy or protein assays. Cells were fixed with 4% paraformaldehyde for 15 minutes at room temperature and then permeabilized with 0.1% Triton X-100 in PBS for 10 minutes. After washing with PBS, cells were blocked with 1% BSA in PBS for 30 minutes to reduce nonspecific binding. Actin filaments were stained by incubating cells with Phalloidin (Alexa Fluor 488, 1:200 dilution) in 1% BSA for 30 minutes at room temperature. Following a PBS wash, nuclei were stained with DAPI (300 nM) for 5 minutes, cells were covered with antifade solution (Sigma-Aldrich S7114).

### 1.10 GUT-ON-A-CHIP ORGANOID SYSTEM

We created a mesofluidic human intestinal chip with polydimethylsiloxane (Sylgard™ 184, Krayden DC4019862) via a standard soft-lithography approach. The chip consists of three distinct channels: a "lumen channel" representing the intestinal lumen, a "side channel" mimicking the circulatory system, and a "stromal gel guide" channel separating the lumen and side channel. We introduced hydrogel precursor material to the stromal gel guide channel to establish a gel wall structure supported by the ridges extending from the top of each side of the stromal gel guide channel. The gel wall segregated the lumen channel and side channel into two sealed compartments and served as a substrate to support the growth of organoid-derived human intestinal epithelial cells. The gel wall was comprised of cross-linked neutralized type I collagen. Briefly, rat tail type I collagen (5 mg/mL; Cultrex 3-D Culture Matrix, R&D Systems™ 344702001) was diluted with neutralization buffer composed of HEPES (20 mM, Gibco™ 15630080), NaOH (9.4 mM), and NaHCO3 (53 mM, Sigma-Aldrich S5761) in PBS (Fisher BioReagents BP661-10) to prepare 2 mg/mL neutralized type I collagen. The neutralized collagen solution was introduced into the stromal gel guide channel and underwent gelation at 37°C for 1 h. We then cross-linked the collagen gel wall via 1-ethyl-3-(3-dimethylaminopropyl) carbodiimide (Thermo Scientific 22980)–N-hydroxysuccinimide (Sigma-Aldrich 130672) chemistry through the side channel at 4°C for 40 min to strengthen its mechanical properties as previously described[8,9]. The mesofluidic intestinal chip was sterilized one day before cell seeding using 70% ethanol. Briefly, we filled the lumen and side channels with 70% ethanol and immersed the whole device in 70% ethanol for 30 s; we then washed the ethanol from the chip surface with sterile water and rinsed the channels with PBS three times.

Primary human epithelial cells were derived from duodenal organoids (donor H416) obtained from the Harvard Digestive Diseases Center Organoid Core (David Breault Lab) at Boston Children's

Hospital as previously described[10]. Briefly, human duodenal stem cells obtained by disrupting organoids were suspended in expansion media at a density of 1.43 x $10^6$ cells/mL (or 1.42×106 cells/$cm^2$ with respect to the surface area of the collagen gel wall), and 75 µL of cell suspension was introduced into the lumen channel. The expansion media consisted of 50% vol/vol advanced DMEM/F12 medium (Gibco™ 12634010) and 50% vol/vol WRN conditioned medium (Harvard Digestive Diseases Center Organoid Core, David Breault Lab) supplemented with 1x GlutaMax (Gibco™ 45050061), 10 mM HEPES (Gibco™ 15630080), 1 mM N-acetyl-cysteine (Sigma-Aldrich A8199), 10 mM nicotinamide (Sigma-Aldrich N0636), 1x B27 (Gibco™ 12587010), 1x N2 (Gibco™ 17502048), 10 nM gastrin (Sigma-Aldrich G9145), 500 nM A83-01 (Sigma-Aldrich SML0788), 50 ng/mL murine epidermal growth factor (PeproTech 315-09), 3 µg/mL SB202190 (Sigma-Aldrich S7067), 10 µM Rho-kinase inhibitor (Y-27632, Sigma-Aldrich Y0503), and 50 µg/mL Primocin (InvivoGen ant-pm-2). We maintained the chips in a cell culture incubator at 37°C tilted at a 90° angle, with the lumen channel facing up throughout the cell culture period, allowing cells to settle onto the collagen gel wall to form the epithelium. The chips were fed daily with fresh expansion media for 3–5 days and then differentiated via differentiation media under air–liquid interface conditions (media in the side channel and air in the lumen channel) as previously described[11] for 5 days before the introduction of NPs to the lumen channel. Differentiation media consisted of 80% vol/vol advanced DMEM/F12 medium and 20% R-spondin conditioned medium (Harvard Digestive Diseases Center Organoid Core, David Breault Lab) with the same supplements as the expansion media, with the exception of nicotinamide and SB202190 and the addition of 100 ng/mL Noggin (PeproTech 250-38), 10 µM N-[N-(3,5-difluorophenacetyl-L-alanyl)]-(S)-phenylglycine t-butyl ester (Sigma-Aldrich 565770), and 1.4 nM prostaglandin E2 (Tocris 2296).

At the conclusion of the 5-day differentiation period, medium was removed from the lumen channel of the chip and replaced with 5 µg/mL Hoechst 33342 (Thermo Scientific™ 62249) and

10 µg/mL wheat germ agglutinin conjugated with Alexa Fluor 647 (Invitrogen™ W3246) in differentiation media to stain nuclei and mucin sugars, respectively. After 30 min of incubation at 37°C, 50 µL of NPs (at ~0.1 mg/mL NP concentrations) was spiked into the lumen channel and imaged for approximately 30 min via confocal microscopy within the chip.

### Supplementary Figures

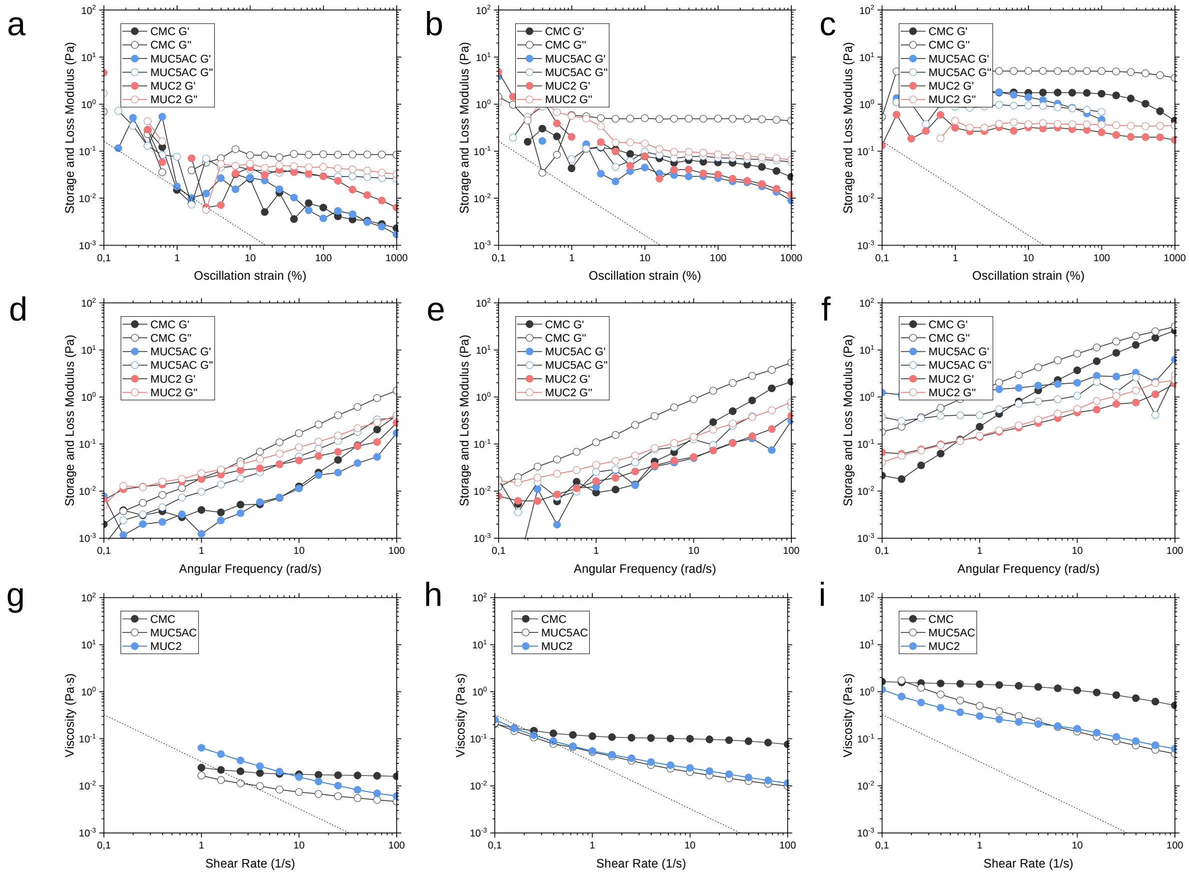


**Supplementary Figure S1: Rheological measurements of gels**. Strain-sweep comparison of (a) 0.5, (b) 1.0, and (c) 2 wt% polymer concentrations in 50 mM HEPES, pH 6.5, 100 mM NaCl. Storage and loss modulus comparison of (d) 0.5, (e) 1.0, and (f) 2 wt% polymer concentrations in 50 mM HEPES, pH 6.5, 100 mM NaCl. Viscosity comparison of (g) 0.5, (h) 1.0, and (i) 2 wt% polymer concentrations in 50 mM HEPES, pH 6.5, 100 mM NaCl. Dashed lines show the nominal sensitivity limits of the rheometer.

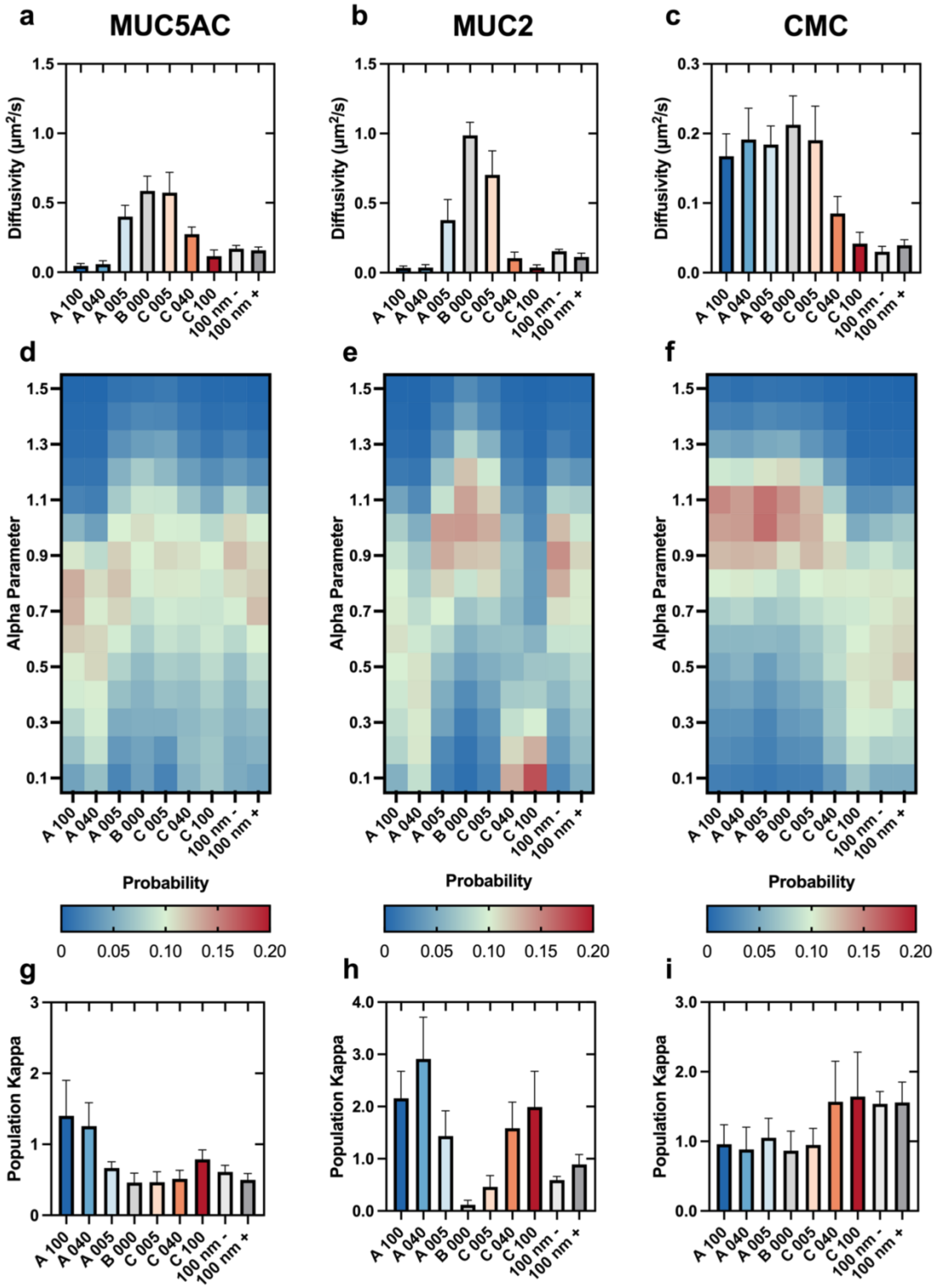


**Supplementary Figure S2: Diffusivity of NPs in 1 wt% polymer solutions.**

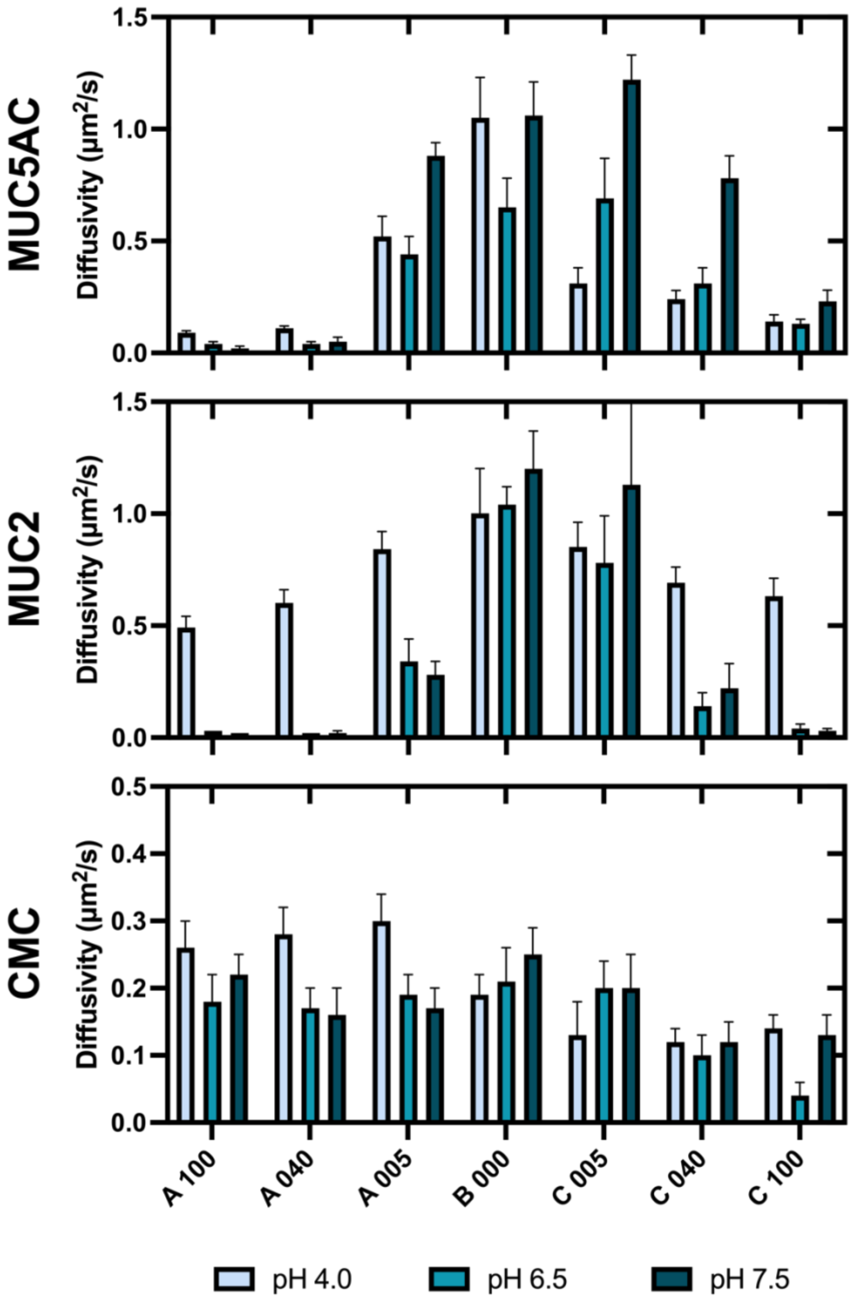


**Supplementary Figure S3: Diffusivity of NPs in various polymer solutions at different pH values.** SPT measurements of particle diffusivity in 1 wt% MUC5AC, MUC2, and CMC at different pH values. Diffusivity calculated from 1 s.

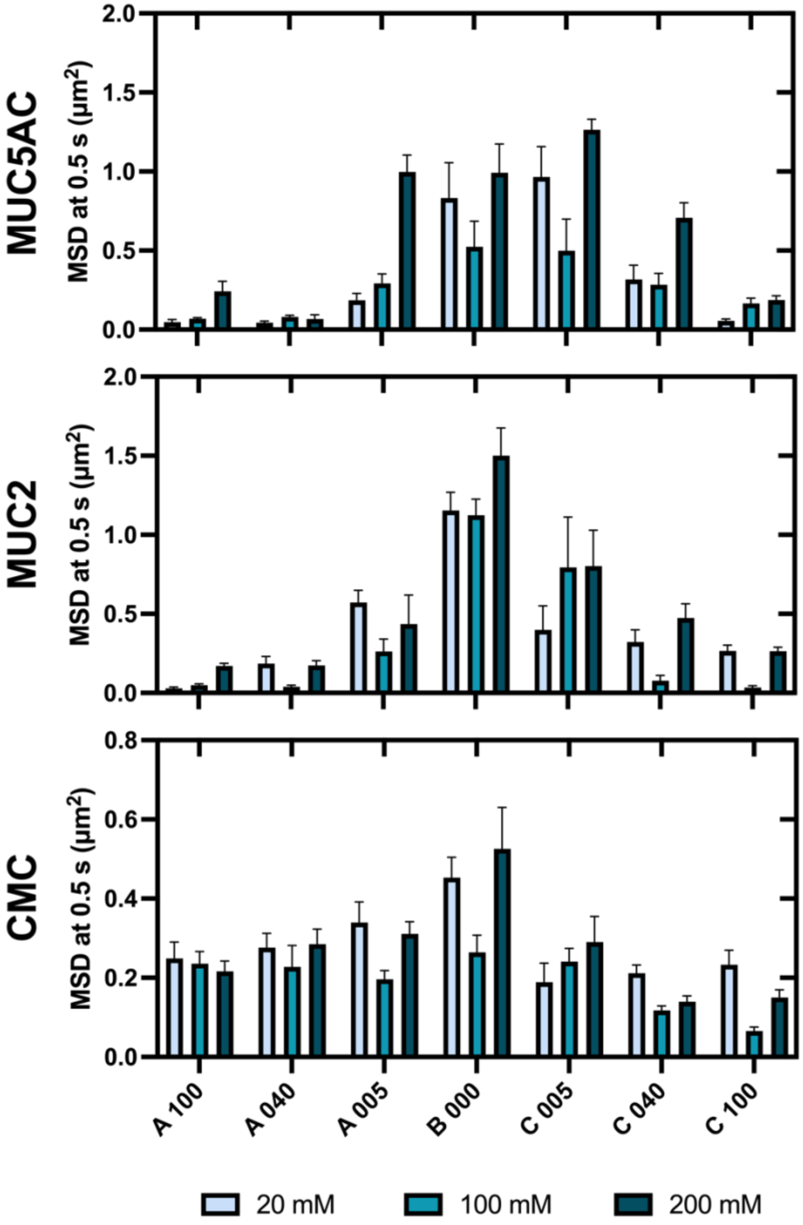


**Supplementary Figure S4: Diffusivity of NPs in various polymer solutions at different ionic strengths.** SPT measurements of particle diffusivity in 1 wt% MUC5AC, MUC2, and CMC in solutions with different ionic strengths. Diffusivity calculated from 1 s.

### Supplementary Tables

#### Supplementary Table S1: Zeta potential values for negatively (COOH) and positively ($NH_3$) charged polystyrene particles pre- and post-MUC2 titration.

| | Pre-MUC2 | | Post-MUC2 | |
|---|---|---|---|---|
| Particle | Zeta (mV) | Standard (mV) | Zeta (mV) | Standard (mV) |
| COOH - 1000 | -67.7 | 10.2 | -64.1 | 7.02 |
| COOH - 500 | -47.7 | 9.8 | -52.8 | 6.23 |
| COOH - 200 | -62.3 | 5.87 | -57.3 | 5.92 |
| COOH - 100 | -64.6 | 8.8 | -54.1 | 5.08 |
| NH3 - 1000 | 35.6 | 11 | -24.5 | 8.17 |
| NH3 - 500 | 52.9 | 6.18 | 3.3 | 7.38 |
| NH3 - 200 | 56.3 | 6.77 | -9.38 | 3.88 |
| NH3 - 100 | 59.8 | 6.41 | 2.7 | 7.11 |

#### Supplementary Table S2: Zeta potential values for NPs at different pH values.

| | pH | | | | | |
|---|---|---|---|---|---|---|
| | 4.0 | | 6.5 | | 7.5 | |
| | Zeta (mV) | Standard (mV) | Zeta (mV) | Standard (mV) | Zeta (mV) | Standard (mV) |
| A 100 | -32.46 | 1.30 | -38.03 | 0.19 | -36.42 | 1.29 |
| A 040 | -28.09 | 0.89 | -34.27 | 0.77 | -34.16 | 1.22 |
| A 005 | -6.20 | 0.94 | -9.41 | 1.42 | -8.04 | 1.87 |
| B 000 | -4.41 | 0.08 | -4.25 | 0.20 | -4.02 | 0.65 |
| C 005 | 1.24 | 1.16 | 1.93 | 0.29 | -1.91 | 1.71 |
| C 040 | 0.80 | 0.42 | 16.22 | 1.77 | 1.30 | 0.67 |
| C 100 | 13.57 | 0.04 | 29.86 | 0.37 | 19.94 | 0.55 |

#### Supplementary Table S3: Zeta potential values for NPs at different ionic strengths.

| | Ionic strength | | | | | |
|---|---|---|---|---|---|---|
| | 20 mM | | 100 mM | | 200 mM | |
| | Zeta (mV) | Standard (mV) | Zeta (mV) | Standard (mV) | Zeta (mV) | Standard (mV) |
| A 100 | -42.90 | 0.46 | -38.03 | 0.19 | -32.39 | 0.88 |
| A 040 | -39.02 | 0.59 | -34.27 | 0.77 | -28.81 | 1.91 |
| A 005 | -14.39 | 0.18 | -9.41 | 1.42 | -7.63 | 0.74 |
| B 000 | -8.85 | 1.53 | -4.25 | 0.20 | -4.56 | 0.16 |
| C 005 | 2.89 | 0.64 | 1.93 | 0.29 | 2.15 | 0.75 |
| C 040 | 24.50 | 1.45 | 16.22 | 1.77 | 13.40 | 1.64 |
| C 100 | 33.51 | 1.49 | 29.86 | 0.37 | 23.41 | 1.32 |
